# InfoTrim: A DNA Read Quality Trimmer Using Entropy

**DOI:** 10.1101/201442

**Authors:** Jacob Porter, Liqing Zhang

## Abstract

Biological DNA reads are often trimmed before mapping, genome assembly, and other tasks to improve the quality of the results. Biological sequence complexity relates to alignment quality as low complexity regions can align poorly. There are many read trimmers, but many do not use sequence complexity for trimming. Alignment of reads generated from whole genome bisulfite sequencing is especially challenging since bisulfite treated reads tend to reduce sequence complexity. InfoTrim, a new read trimmer, was created to explore these issues. It is evaluated against five other trimmers using four read mappers on real and simulated bisulfite treated DNA data. InfoTrim produces reasonable results consistent with other trimmers.

## 1. Introduction

Raw DNA reads generated by next generation sequencing machines can have diminishing quality on the 5’ ends [1], [2] and be contaminated by adapters [3]. These errors can affect the quality of alignment and downstream SNP, indel, and mehtylation calling [3]. Read trimming is a process where the raw DNA reads are cut to address issues of low quality or adapter contamination.

This study introduces a new read quality trimmer called InfoTrim derived from the maximum information approach of the popular trimmer Trimmomatic [4]. InfoTrim introduces an entropy term, a measure of sequence complexity, to the maximum information model of Trimmomatic. This term was added since sequence complexity can correlate with alignment quality as low complexity reads are more frequently unmapped or mapped to multiple locations while high complexity reads are more frequently uniquely mapped [5]. InfoTrim uses Shannon entropy, which measures sequence complexity by counting the frequency of bases in a read. Shannon entropy plays a fundamental role in information theory [6].

Some other read processing tools incorporate sequence complexity, but they don’t use Shannon entropy in the same way as InfoTrim does. The trimmer Reaper uses a measure of tri-nucleotide complexity that is the same as is used in the base masking program DUST [7], [8]. The read mapper Novoalign trims low complexity 5’ tails up to 5-9 bp [9]. The bisulfite DNA read mapper BatMeth filters out reads with low Shannon entropy [10]. The UEA Toolkit has a low sequence complexity filter that eliminates reads with at most two bases, but it is for RNA reads [11].

The other terms in the InfoTrim model include the sequence length and the phred score. There is a term for sequence length threshold and for the total sequence length. Everything else being equal, preserving the length of a read is generally preferred as this uses more of the raw DNA read data. Sequence length can affect indel calling [12]. The phred score of a DNA read gives a per-base measure of the probability that the base was called correctly [13]; thus, bases near the 5’ end of the read with low phred score can be trimmed. These terms are the same as the Trimmomatic model, but InfoTrim applies the geometric mean to the terms so that the input parameters can be interpreted more intuitively. The geometric mean is a balanced mean that does not depend on the scale of the terms. InfoTrim does not support adapter trimming at this time, but there are many trimmers for this including Trimmomatic.

To compare the performance of InfoTrim, data involving bisulfite treated DNA reads (BS-Seq) was used. Bisulfite treatment of DNA converts unmethylated cytosine to thymine upon PCR amplification while leaving methylated cytosine unchanged. It is a way to study epigenetic methylation, which relates to disease and development [14]. BS-Seq data was used since aligning bisulfite-treated DNA is a challenging task since bisulfite treatment tends to reduce sequence complexity [5] and can introduce major biases [15].

This study compares InfoTrim with five other trimmers involving four read mappers. The read trimmers were Cutadapt [16], Erne-filter [3], Reaper [7], Sickle [17], and Trimmomatic [4]. The read mappers were BisPin [18], Bismark [19], BWAmeth [20], and Walt [21]. Table 1 gives information on the read trimmers used in this study.

**TABLE 1:** Trimmer features, information, and settings used in this study

| Trimmer | Version | Website | Algorithm Type | Language | Features | Settings |
| --- | --- | --- | --- | --- | --- | --- |
| InfoTrim | 0.1.0 | github.com/<br>JacobPorter/InfoTrim | Running Score | Python 2.7 | has an entropy term,<br>multiprocessing | s:0.1, r:0.4, m:25, t:0 |
| Trimmomatic | 0.36.0 | www.usadellab.org/<br>cms/?page=trimmomatic | Running Score,<br>Windowed | Java | Adapter stripping,<br>multithreaded | MAXINFO: 25: 0.5 |
| Cutadapt | 1.14 | cutadapt.<br>readthedocs.io/<br>en/stable/guide.html | Running Score | C and Python | Adapter stripping | -q 10 |
| Erne-filter | 2.1.1 | erne.sourceforge.net/ | Running Score | C++ | contaminant removal,<br>multithreaded | Default |
| Reaper | 13.100 | www.ebi.ac.uk/<br>stijn/reaper/reaper.html | Quality Score | C | demultiplexes<br>barcoded sequences,<br>adapter stripping,<br>tri-nucleotide<br>complexity score | -geom no-bc -3pa ""<br>-tabu "" -dust-suffix<br>90 -qqq-check 50/15<br>-nozip |
| Sickle | 1.33 | github.com/<br>najoshi/sickle | Windowed | C | Supports gzip files | -t sanger |

## 2. Implementation

InfoTrim is a multiprocess Python 2.7 program featuring a maximum information score used to cut DNA reads at the 5’ end. For a read *d*, the substring *d*_*l*_ is the read starting at base position 0 and ending at base position *l*. Starting at the first base, the score *S*_*total*_(*d*_*l*_) is computed for all *l* starting at 0 and ending at the total length of the read *d*. The position *l* that maximizes the score is chosen and every base after *l* is discarded from the read. The InfoTrim maximum information model involves four terms.

The first term, *S*_*len*_(*d*_*l*_), is a logistic function that provides a length threshold *m*. This causes the trimmer to strongly prefer reads larger than *m*. The reason for this is that reads that are too small will have too little information to be useful. The default is *m* = 25, and the equation is the following.

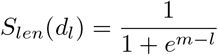

The second term, *S*_*cov*_ (*d*_*l*_), causes the trimmer to prefer longer reads to shorter reads in order to increase the coverage that the read represents. It is simply the length of the read, giving a linear relationship between the length and the score. It is described in the following.

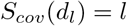

The phred score, *S*_*phred*_(*d*_*l*_), is the third term, and it measures base calling correctness. The phred score at base position *i* in read *d* is the probability that the base was called correctly as determined by the sequencing platform. The phred number, a positive integer, at base position *i* is given by *Q*_*i*_. The probability derived from *Q*_*i*_ is given by 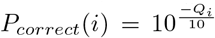. The phred term is given by the following formula. The rationale for using the product is given in the Trimmomatic paper [4].

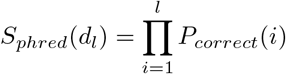

InfoTrim adds a fourth term for entropy. The probability *P*_*l*_(*b*) is the frequency that base *b* is found in read *d*_*l*_. The Shannon entropy is multiplied by one half to give it a scaling between 0 and 1. The formula that InfoTrim uses is given as the following.

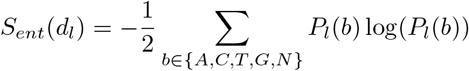

The last three terms are combined using the geometric mean so that proportional changes in each term will contribute equally to the final score. Parameters *r* (for entropy) and *s* (for phred) are used to weight the relative contributions of the last three terms. The values *r*, *s*, and 1 − *r* − *s* must all add up to one. For example, if *r* = 1, then only the entropy term will be used while the coverage and phred scores will not be used. Thus, the final score is the following.

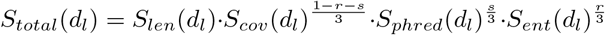

## 3. Data Analysis Methods

Real and simulated data was used to assess the accuracy of InfoTrim. The arguments used for each trimmer are given in Table 1. Default settings were used when available. Otherwise, reasonable settings were used. The DUST score was used with Reaper to compare its sequence complexity term with InfoTrim. For the read mappers, BisPin 0.1.1, Walt 1.0, Bismark 0.16.3, and BWAmeth 0.2.0, default settings were used. A single BisPin index with mask “11111111100111111111” was generated. The FASTQ reads files were first trimmed with the read trimmers and then mapped with the read mappers.

### 3.1. Simulated Data

Five hundred thousand simulated 150bp Illumina bisulfite reads were generated with Sherman using the mouse reference genome (GRCm38.p5) [22] and a two percent error rate, a twenty percent CG context conversion rate, and a two percent CH conversion rate. These settings were chosen since they are realistic. The error rate simulates both sequencing error and natural SNP variation. Sherman does not simulate indels and has a flat phred score. Five hundred thousand simulated Ion Torrent 200bp reads were generated with DWGSIM using the human reference genome (GRCh38.p9) [23]. For DWGSIM, the bisulfite treatment was simulated on the reference genome with the CpG rate as 0.215, the CH rate 0.995, and the over conversion and under conversion rates of 0.0025. This was done with custom Python scripts provided by another lab member. DWGSIM generates paired end data, but single end data was desired, so appropriate ends were chosen to simulate single end data. DWGSIM was used with the following realistic settings: “dwgsim -e 0.012 -E 0.012 -d 250 -s 30 -S 0 -N 1000000 -c 2 -1 200 -2 200 -f TACGTACGTCTGAGCATCGATCGATGTACAGC.” This data was used to compare InfoTrim with other trimmers. DWGSIM simulates the overcalling and undercalling of homopolymer runs that are characteristic of Ion Torrent reads [2]. This higher error rate makes Ion Torrent reads more challenging to align.

Another data set of 500k 100bp simulated Illumina human DWGSIM reads were used to explore the effect of varying the *r* and *s* parameters for InfoTrim. Data was generated with settings that were the same as for the Ion Torrent reads except that the “-f” parameter was not used, and 100bp reads were generated. Varying the *r* and *s* parameters was done with a step size of 0.1. All the read mappers were used to align and map the data.

Recall, the percentage of the total reads aligned correctly, and precision, the percentage of uniquely mapped reads aligned correctly, was calculated for each alignment SAM file generated by the read mappers. These statistics control for false positives and false negatives. A read was correctly aligned if it was mapped to within 3bp of the true location’s starting position. The F1-score, the harmonic mean of precision and recall, was calculated to give a single numeric score for each trimmer and mapper pipeline. A higher F1-score indicates better accuracy performance.

The mapper Walt couldn’t always map the output from Reaper, and some pipelines and data had zero for Reaper results. For this reason, Walt or Reaper were sometimes excluded from the results.

### 3.2. Real Data

Five hundred thousand real whole mouse genome bisulfite 101 bp Illumina HiSeq 2000 reads were downloaded from the SRA trace archive with accession number SRR921759. The data was trimmed with the read quality trimmers and then mapped with BisPin. A strict alignment quality filter of 85 was used since reads passing this filter should have a precision approaching one hundred percent, so the unique alignments are presumptively correct. A filter value of 96 indicates that the read perfectly matches the genome, so a filter value of 85 indicates very high quality alignments.

A second real data set consisting of one million hairpin bisulfite-treated Illumina mouse reads was taken from the data described in [14], and this hairpin data was used to compare pipelines using presumptively correct alignments in the following manner. Hairpin bisulfite data uses paired-end sequencing such that the original DNA strand, untreated by bisulfite, can be recovered [18]. The original strand generally has more entropy, and can align better than bisulfite treated strands [5]. This recovery procedure was performed using BisPin, which recovered approximately 600k reads, and the resulting original reads were aligned with BFAST [24], Bowtie2 [25], and BWA [26] to create three files of presumptively correct unique alignments. The corresponding one million C-to-T converted bisulfite forward direction reads were then trimmed and aligned with all of the tools used in this study. The uniquely mapped reads from each trimmer and bisulfite mapper pipeline were then compared to the presumptively correct reads of each regular read mapper and averaged.

## 4. Results and Discussion

### 4.1. Simulated Data

Figure 1 shows the results of varying the *r* and *s* parameters on F1-score for each read mapper. The left-most pane of Figure 1 shows the F1-score when no trimming was performed. The highest F1-score and (*r*, *s*) arguments for each mapper are the following: Bismark 0.910 (0.3, 0.1), BisPin 0.881 (0.4, 0.1), BWAmeth 0.880 (0.1, 0.0), and Walt 0.914 (0.1, 0.2). This setting allowed BWAmeth to beat the untrimmed F1-score by 7.40E-5, a small amount. The other mappers had similar performance. According to Figure 1, low to moderate settings for *r* and low settings for *s* substantially outperformed other settings. This suggests that the coverage term, *S*_*cov*_(*d*_*l*_), should have a high weight and is more important than entropy and the phred score; however, trimming with entropy can still usefully improve the accuracy. For this reason, *r* = 0.4 and *s* = 0.1 were chosen as the default settings for InfoTrim. Bismark and BisPin were generally the most consistent with the F1-score falling the least in the right-most panes. The quality of Walt and BWAmeth fell off more quickly.

**Figure 1:**
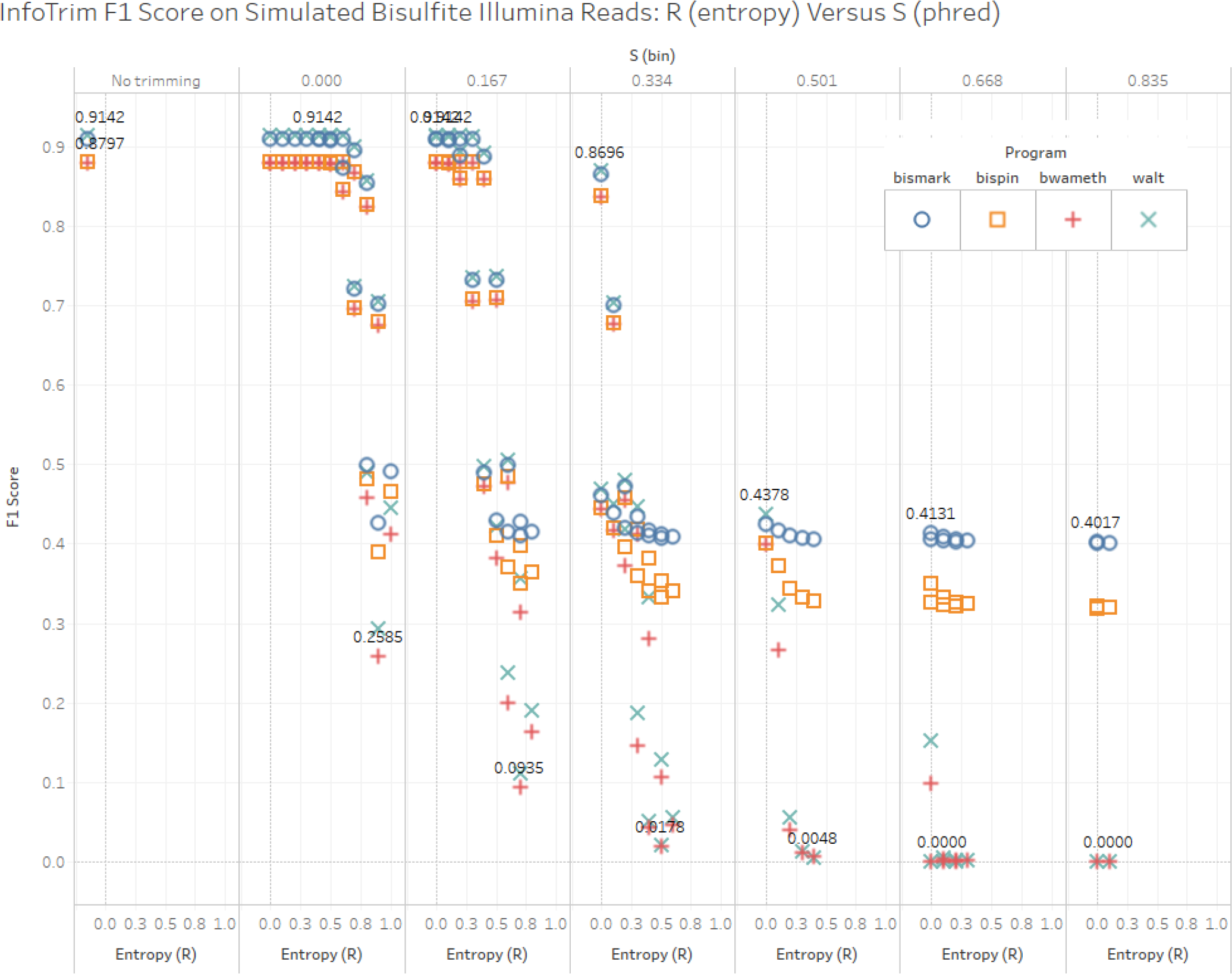
InfoTrim simulation results vis-a-vis systematically varying the r and s parameters.

On the Sherman simulated Illumina mouse reads, InfoTrim improves the F1-score better than all the other trimmers except for pipelines involving BWAmeth as shown in Figure 2. With the BWAmeth pipeline, InfoTrim was second best beating Reaper, the other trimmer that uses sequence complexity, and it beat Trimmomatic, the trimmer that InfoTrim derives from. Pipelines involving InfoTrim don’t always have the highest percentage of uniquely mapped reads, but more uniquely mapped reads don’t always equate to a higher accuracy as the F1-score shows.

**Figure 2:**
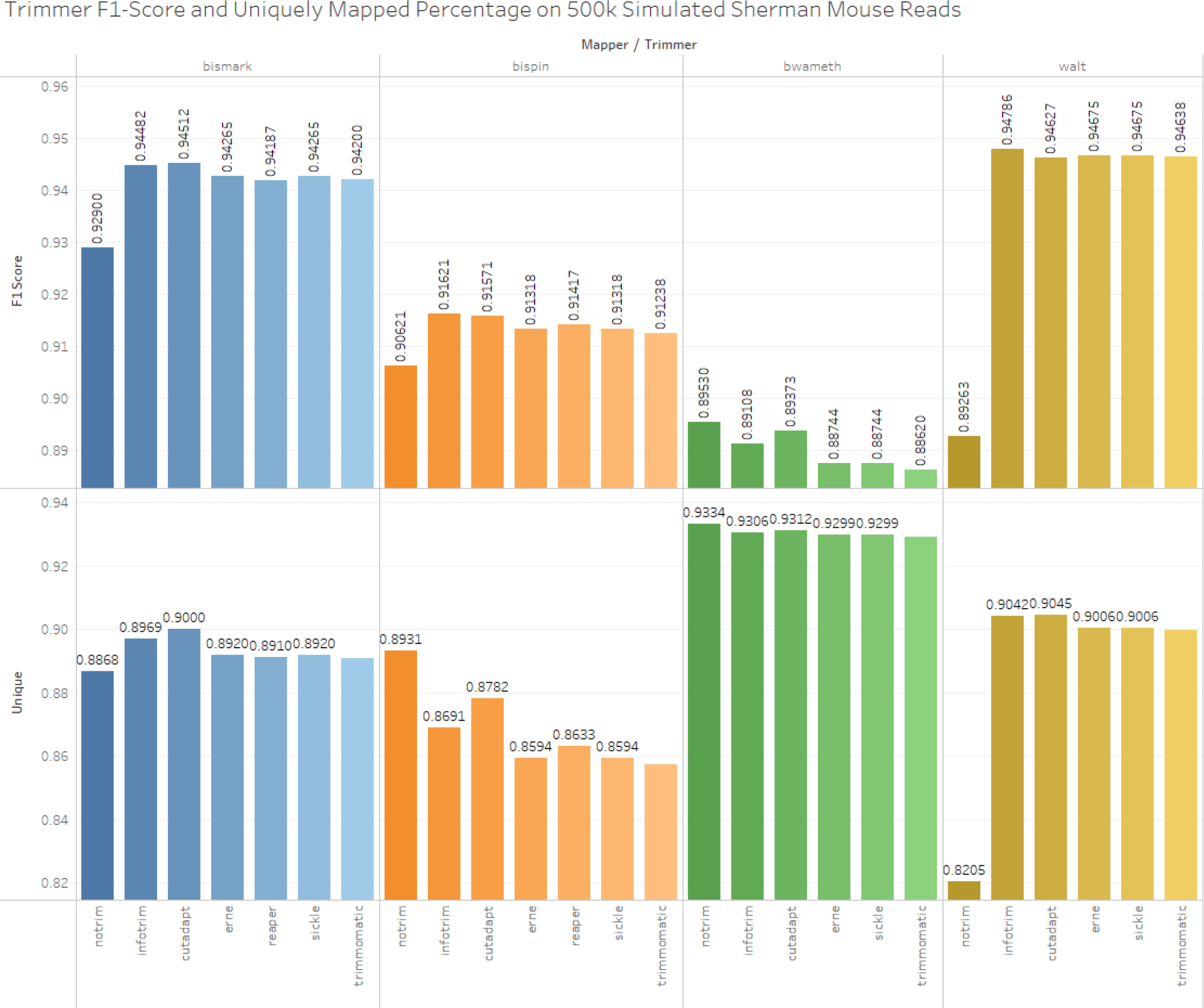
Sherman mouse Illumina simulation on 500k reads with F1-score and uniquely mapped percentage

Figure 3 shows a plot of precision versus recall while the numeric label is the F1-score on the DWGSIM simulated human Ion Torrent bisulfite treated reads. InfoTrim, Cutadapt, and Reaper performed similarily while Trimmomatic was the next best. Erne-filter and Sickle were the worst performing. The read mapper BisPin generally had the highest F1-score for all pipelines with the exception of Erne-filter and Sickle. However, trimming did little to improve the F1-score compared to no trimming.

**Figure 3:**
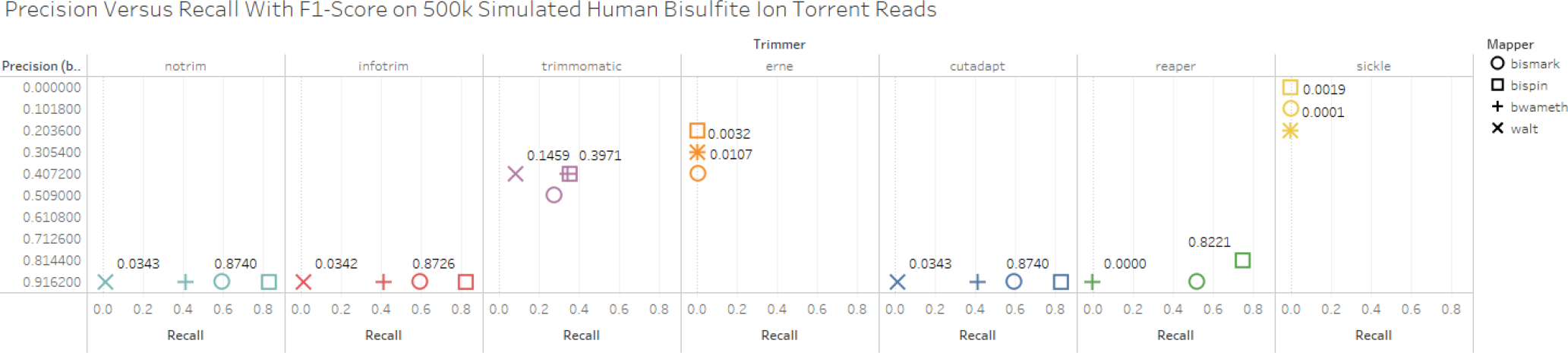
Precision versus recall and numeric F1-score on DWGSIM human Ion Torrent simulated bisulfite reads

### 4.2. Real Data

On the strict filter test of real mouse reads, all trimmers improved the alignment score as indicated in Figure 4. InfoTrim performed the best on the real data and aligned 1300 more reads than Cutadapt, the closest competitor. InfoTrim aligned 15,025 more reads than without trimming.

**Figure 4:**
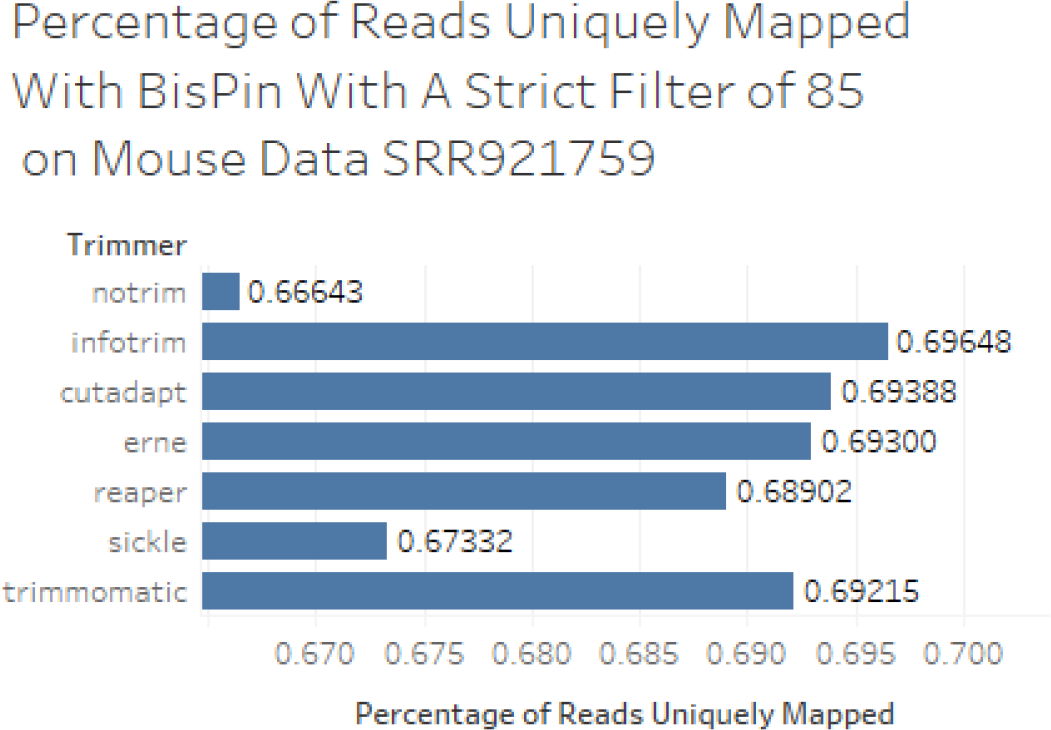
Percentage of 500k mouse real reads aligned presumptively correctly with a strict BisPin filter of 85.

The proportion of presumptively correct alignments for the hairpin data by trimmer and mapper is shown in Figure 5. According to the average proportion of reads aligned correctly across all mappers, InfoTrim was third best overall, behind Cutadapt and Erne-filter; however, there is very little difference in this score except for Reaper, which did poorly since BWAmeth and Walt did not map this data well. Pipelines invovling BisPin were generally good, but BWAmeth often did better except for the pipeline with the trimmer Reaper. Pipelines with Bismark were the worst. This shows that the results for InfoTrim are reasonable and consistent with other trimmers.

**Figure 5:**
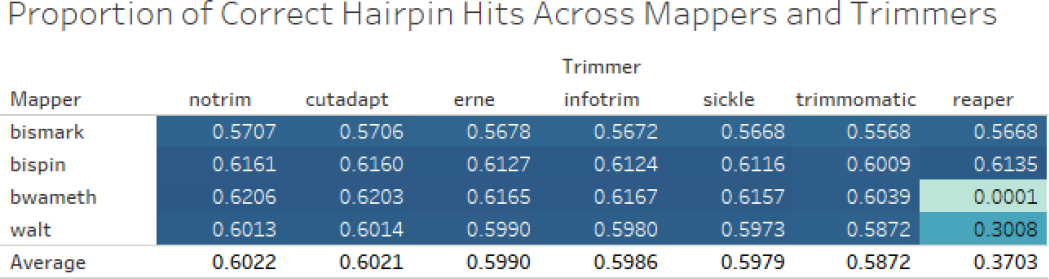
Hairpin data accuracy analysis. The cell for a trimmer and mapper represent an average of proportions of correct alignments using presumptively correct alignments from BFAST, BWA, and Bowtie2 on recovered reads. The average performance across all bisulfite mappers for a given trimmer is shown in the row “Average.”

### 4.3. Timing

A run time analysis was performed by trimming 5 million real human bisulfite reads consisting of 2 million reads from SRR1104850, 2 million reads from SRR1799718, and one million reads from SRR3106764. All the read trimmers except for InfoTrim took approximately one minute to complete using a single thread. Trimmomatic was the fastest with 30s and Reaper was the slowest with 1m21s. InfoTrim was considerably slower and took 72m49s with a single process and 15m26s with six processes, an improvement of 4.7 times. InfoTrim is probably slow since it is implemented in Python. Timing was calculated with the linux time command, and the read trimmers were run on the Virginia Tech CS Department’s bioinformatics machine mnemosyne2 consisting of 16 processing cores of Intel(R) Xeon(R) CPU E5620 @ 2.40GHz with 132 GB of memory.

## 5. Conclusion

Using sequence complexity to improve read trimming is a promising strategy as sequence complexity can relate to alignment accuracy performance. InfoTrim is a new Python read trimmer that performs better than five other trimmers on simulated Sherman bisulfite data and real mouse data involving alignment pipelines of four read mappers. InfoTrim did not substantially reduce accuracy on simulated human Ion Torrent bisulfite reads unlike some read trimmers. Unfortunately, InfoTrim is slow, but the techniques could be incorporated into the Java based Trimmomatic, the fastest read trimmer. The only other read trimmer studied that used sequence complexity was Reaper, and InfoTrim outperformed this trimmer. Given the presented results, perhaps sequence complexity should be incorporated into more read trimmers.

## Acknowledgments

Jacob Porter created InfoTrim and wrote the paper, and Liqing Zhang consulted. Hong Tran provided the bisulfite simulation software used to prepare the reference genomes for DWGSIM. There are no conflicts of interest to declare.

